# Metatranscriptomic and thermodynamic insights into medium-chain fatty acid production using an anaerobic microbiome

**DOI:** 10.1101/424762

**Authors:** Matthew J. Scarborough, Christopher E. Lawson, Joshua J. Hamilton, Timothy J. Donohue, Daniel R. Noguera

## Abstract

Biomanufacturing from renewable feedstocks represents a sustainable strategy to offset fossil fuel-based chemical production. One potential biomanufacturing strategy is production of medium chain fatty acids (MCFA) from organic feedstocks using either pure cultures or microbiomes. While the set of microbes in a microbiome can often metabolize more diverse organic materials than a single species, and the role of specific species may be known, knowledge of the carbon and energy flow within and between organisms in MCFA producing microbiomes is only starting to emerge. Here, we integrate metagenomic, metatranscriptomic, and thermodynamic analyses to predict and characterize the metabolic network of an anaerobic microbiome producing MCFA from organic matter derived from lignocellulosic ethanol fermentation conversion residue. A total of 37 high quality (>80% complete, < 10% contamination) metagenome-assembled genomes (MAGs) were recovered from the microbiome and metabolic reconstruction of the 10 most abundant MAGs was performed. Metabolic reconstruction combined with metatranscriptomic analysis predicted that organisms affiliated with *Lactobacillus* and Coriobacteriaceae degraded carbohydrates and fermented sugars to lactate and acetate. Lachnospiraceae and Eubacteriaceae affiliated organisms were predicted to transform these fermentation products to MCFA. Thermodynamic analyses identified conditions in which H_2_ is expected to be either produced or consumed, suggesting a potential role of H_2_ partial pressure on MCFA production. From an integrated systems analysis perspective, we propose that MCFA production could be improved if microbiomes are engineered to use homofermentative instead of heterofermentative *Lactobacillus*, and if MCFA-producing organisms are engineered to preferentially use a thioesterase instead of a CoA transferase as the terminal enzyme in reverse β-oxidation.

**Importance:** Mixed communities of microbes play important roles in health, the environment, agriculture, and biotechnology. While tapping the combined activity of organisms within microbiomes may allow the utilization of a wider range of substrate over the use of pure cultures for biomanufacturing, harnessing the metabolism of these mixed cultures remains a major challenge. Here, we predict metabolic functions of bacteria in a microbiome that produces medium-chain fatty acids from a renewable feedstock. Our findings lay the foundation to begin addressing how to engineer and control microbiomes for improved biomanufacturing; to build synthetic mixtures of microbes that produce valuable chemicals from renewable resources; and to better understand microbial communities that contribute to health, agriculture, and the environment.

## Introduction

Biological production of chemicals from renewable resources is an important step to reduce societal dependence on fossil fuels. One approach that shows potential for the biological production of chemicals from renewable resources, the carboxylate platform (1, 2), uses anaerobic microbial communities to biotransform complex substrates into carboxylic acids, including medium-chain fatty acids (MCFA). MCFA such as hexanoate (a six carbon monocarboxylate, C6) and octanoate (an eight carbon monocarboxylate, C8) are used in large quantities for the production of pharmaceuticals, antimicrobials, and industrial materials, and can be processed to chemicals currently derived from fossil fuels (3, 4).

Previous applications of the carboxylate platform have focused on converting organics from undistilled corn beer (5, 6), food (7, 8), winery residue (9), thin stillage from corn ethanol production (10), and lignocellulose-derived materials (11–13) to MCFA, and as we have shown for lignocellulosic biofuel production (4), one can anticipate economic benefits of converting organic residues from these industries into MCFA.

MCFA producing bioreactors contain diverse microbial communities (4, 5, 12). While the roles of some community members in these microbiomes can be inferred from studies with pure cultures and from phylogenetic relationships (10, 12, 14, 15), detailed knowledge of specific metabolic activities in many members of these microbiomes is only starting to emerge (16). In general, some community members participate in hydrolysis and fermentation of available organic substrates, while others are involved in the conversion of intermediates to MCFA via reverse β-oxidation, a process also known as chain elongation (1). In reverse β-oxidation, an acyl-CoA unit is combined with acetyl-CoA, with each cycle elongating the resulting carboxylic acid by two carbons(1). Energy conservation in organisms using reverse β-oxidation as the main metabolic process for growth relies on ATP generation with reduced ferredoxin, which is generated through both pyruvate ferredoxin oxidoreductase and an electron bifurcating acyl-CoA dehydrogenase (17). A proton translocating ferredoxin, NAD_ reductase, is used to reduce NAD with ferredoxin and create an ion motive force which is used to generate ATP (17). The even-chain butyric (C4), hexanoic (C6), and octanoic (C8) acids are all potential products of reverse β-oxidation when the process is initiated with acetyl-CoA. The odd-chain valeric (C5) and heptanoic (C7) acids are products of reverse β-oxidation when the chain elongation process starts with propionyl-CoA. While there are demonstrations of this wide range of possible products from chain elongation (5, 18), and MCFA-bioreactors typically produce more than one product (4, 12, 14, 15, 19, 20), a strategy to control the final product length has not yet emerged. We are interested in obtaining the knowledge needed for the rational development and implementation of strategies to improve MCFA yields and control product formation in MCFA-producing microbiomes.

Earlier we reported on a bioreactor that produced a mixture of C2, C4, C6 and C8 from lignocellulosic stillage (4). Based on 16S rRNA tag sequencing, we found that five major genera, three Firmicutes (*Lactobacillus*, *Roseburia*, *Pseudoramibacter*) and two Actinobacteria (*Atopobium*, *Olsenella)*, represented more than 95% of the community (4). Based on the phylogenetic association of these organisms, the *Lactobacillus* and the Actinobacteria were hypothesized to produce lactic acid, while *Roseburia* and *Pseudoramibacter* were hypothesized to produce the even-chain C4, C6, and C8 acids (4). Furthermore, lactic acid was proposed as the key fermentation product that initiated chain elongation in the microbiome (4). However, since phylogenetic association is not enough to understand in detail the metabolism of these organisms, the earlier study did not generate sufficient knowledge to help understand how to control a MCFA-producing microbiome.

Here we report on further studies of the MCFA-producing microbiome reported earlier (4), where we utilized a combination of metagenomic, metatranscriptomic, and thermodynamic analyses to reconstruct the combined metabolic activity of the microbial community. We analyzed the gene expression patterns of the ten most abundant community members during steady-state reactor operation. Our results identify several community members that expressed genes predicted to be involved in complex carbohydrate degradation and in the subsequent fermentation of degradation products to lactate and acetate. Genes encoding enzymes for reverse β-oxidation were expressed by two abundant organisms affiliated with the class Clostridia. Based on a thermodynamic analysis of the proposed MCFA-producing pathways, we predict that individual Clostridial organisms use different substrates for MCFA production (lactate, versus a combination of xylose, H_2_, and acetate). We also show that, under certain conditions, production of MCFA provides energetic benefits compared to production of butyrate, thus generating hypotheses for how to control the final products of chain elongation. This knowledge lays a foundation to begin addressing how to engineer and control MCFA producing microbiomes.

## Results

### Microbiome characterization

We previously described the establishment of a microbiome that produces MCFA in a bioreactor that is continuously fed with the residues from lignocellulosic ethanol production (4). The reactor feed, identified as conversion residue (CR) in Fig. 1, contained high amounts of xylose, carbohydrate oligomers, and uncharacterized organic matter. To gain insight into the microbial activities that were associated with this MCFA-producing microbiome, samples were collected for metagenomic analysis at five different times (Days 12, 48, 84, 96, and 120), and RNA was prepared for metatranscriptomic analysis at Day 96. At the time of metatranscriptomic sampling, the bioreactor converted 16.5% of the organic matter (measured as chemical oxygen demand (COD)) in conversion residue to C6 and C8. During the period of reactor operation described in Fig. 1, the bioreactor converted 16.1 ± 3.1 % of COD to C6 and C8, and, therefore, Day 96 is representative of the overall reactor performance.

**Fig. 1.**
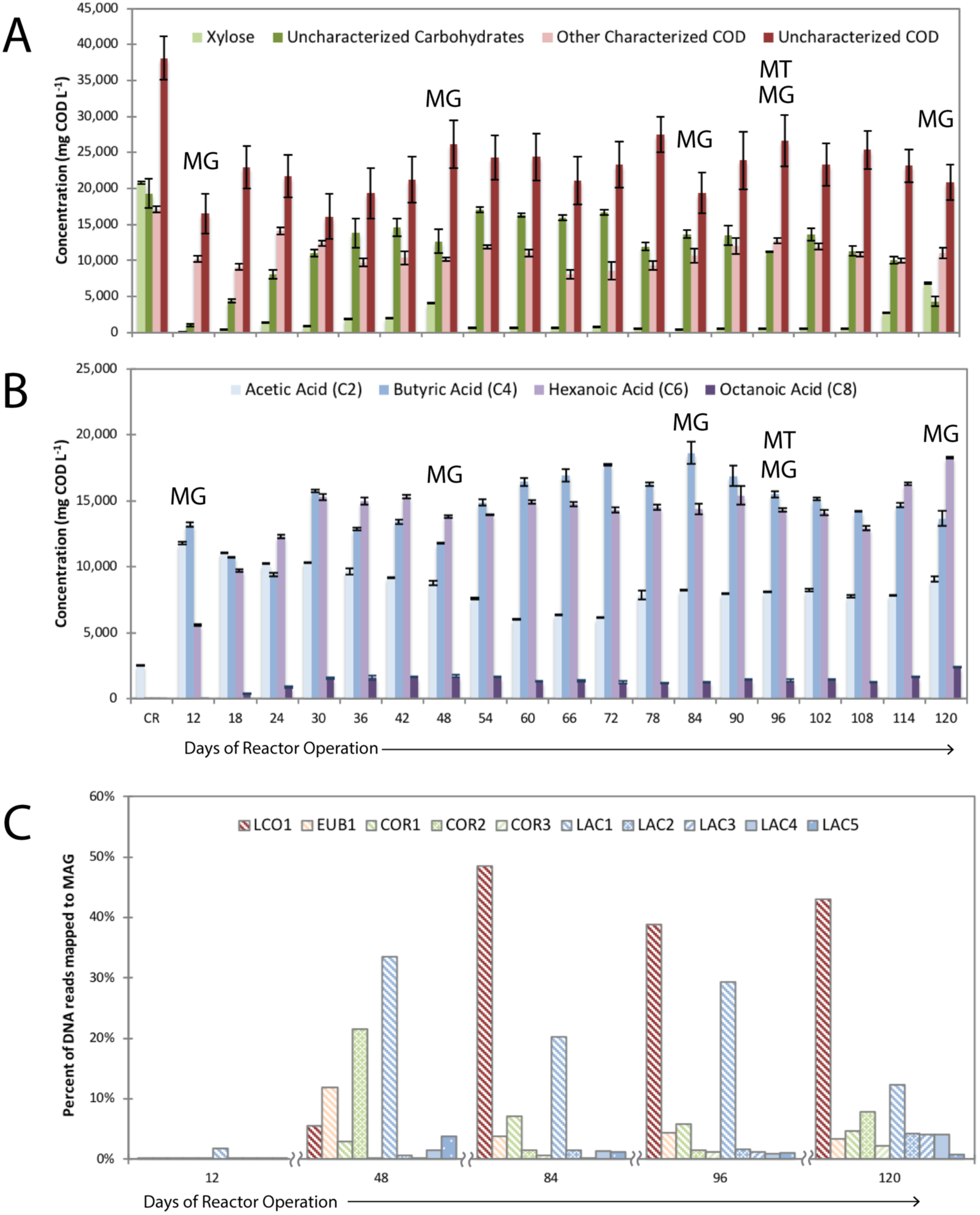
Transformation of materials in lignocellulosic ethanol conversion residue by an anaerobic microbiome and abundance of MAGs. During 120 days of reactor operation, compounds in conversion residue (CR) were converted to medium-chain fatty acids. **For (A) and (B),** the first set of bars in the figure describe concentrations in the feed (CR), whereas the rest of the bars describe concentrations in the reactor. A more detailed description of the operation of this reactor is presented elsewhere (4). Samples were taken for metagenomic (MG) analysis from five timepoints (Day 12, Day 48, Day 84, Day 96, and Day 120 and for metatranscriptomic analysis (MT) from one time point (Day 96). Overall, the bioreactor transformed xylose, uncharacterized carbohydrates and uncharacterized COD to acetic (C2), butyric (C4), hexanoic (C6) and octanoic (C8) acids. The microbial community was enriched in 10 MAGs.

From the metagenomic samples, a total of 219 million DNA reads were assembled and binned, resulting in 37 high quality (>80% complete, <10% contamination) MAGs (Supplement 1). In this study, MAGs are the collection of genes that were assembled into contigs and represent the population of organisms associated with this collection. For the Day 96 sample, 90% of the DNA reads mapped to the ten most abundant MAGs, and each individual MAG mapped to either more than 0.9% of the DNA reads or more than 0.9% of the cDNA reads (Supplement 2).. Abundance of the top 10 MAGs was calculated from the percent of the total DNA reads mapped to the MAGs (Fig. 1c). For the Day 96 sample, relative abundance and expression were compared (Fig. 2; Supplement 2). The most abundant MAGs include a Lachnospiraceae (LCO1, 50%), a *Lactobacillus* (LAC1, 30%), a Coriobacteriacea (COR1, 6.3%), and a Eubacteriaceae (EUB1, 6.0%). Four additional *Lactobacilli* and two additional Coriobacteriacea are also predicted to be within the 10 most abundant MAGs (Fig. 2). The other MAGs (Supplement 1) corresponded to Firmicutes (17 MAGs), Actinobacteria (4 MAGs), Tenericutes (3 MAGs), Bacteroidetes (2 MAGs), and Spirochaetes (1 MAG).

**Fig. 2.**
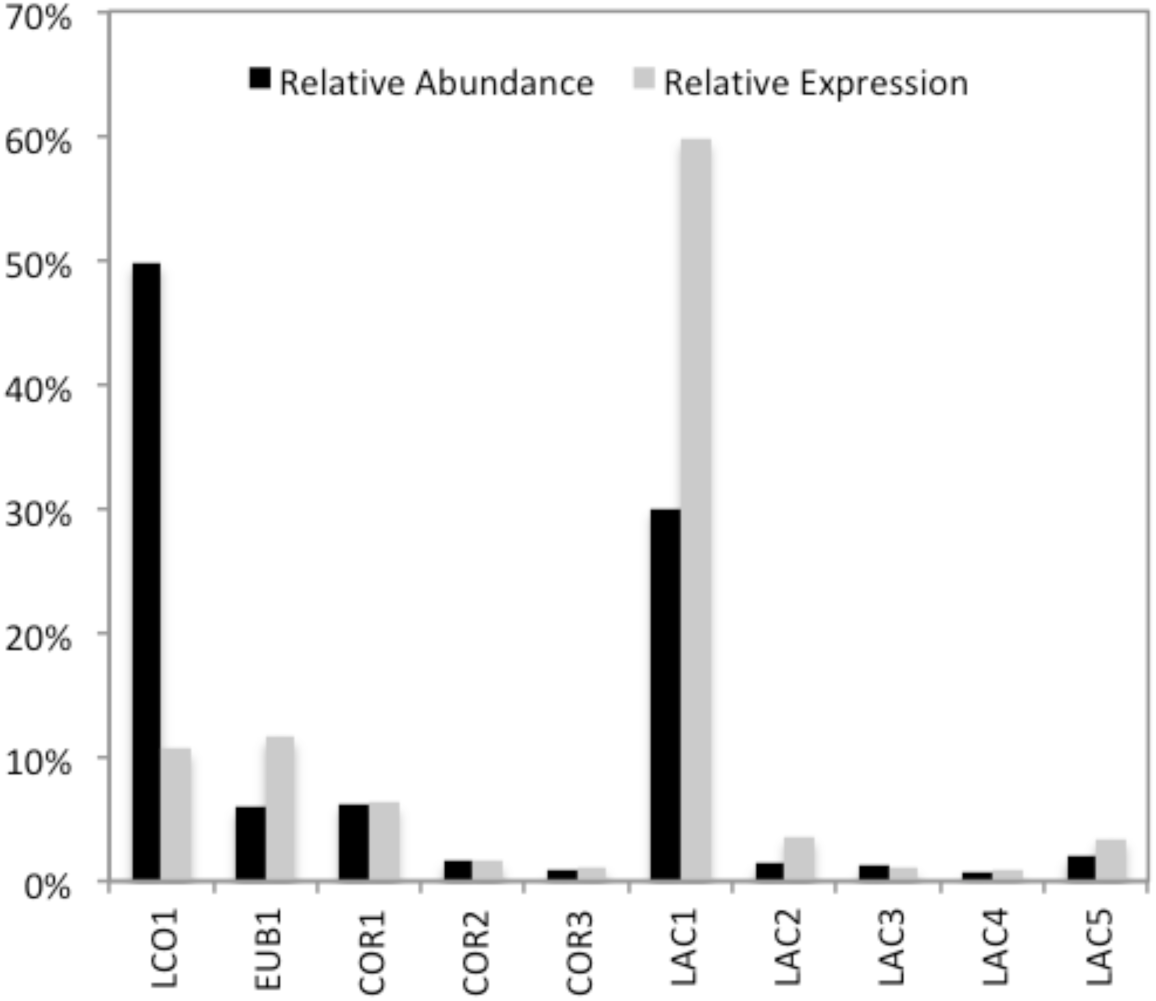
Relative abundance and expression of the 10 most abundant MAGs in the bioreactor at Day 96. Relative abundance was determined by mapping DNA sequencing reads to the MAG and normalizing to the length of the MAG genome. Relative transcript abundance (expression) was determined by mapping c-DNA sequencing reads to the MAG and normalizing to the length of the MAG genome.

The metatranscriptome data, obtained from the Day 96 sample, contained 87 million cDNA reads. After quality checking and removal of rRNA sequences, 82.6 million predicted transcript reads were used for mapping to MAGs. Of these, 85.3% of the predicted transcripts (hereafter referred to as transcripts or mRNA) mapped back to the 10 most abundant MAGs (Supplement 2). Relative expression was calculated from the total filtered mRNA mapped to the MAGs and normalized to the predicted genome length of these bacteria (Fig. 2). MAGs with the highest levels of transcripts included LAC1 (60%), EUB1 (12%), LCO1 (11%), and COR1 (6.3%), which also displayed high abundance in the metagenome (Fig. 2). Whereas LCO1 was most abundant based on DNA reads, LAC1 appeared to have the highest activity based on transcript levels.

A phylogenetic tree of the ten most abundant MAGs was constructed based on concatenated amino acid sequences of 37 single-copy marker genes (Fig. 3). All Bacilli MAGs (LAC1, LAC2, LAC3, LAC4, and LAC5) clustered with *Lactobacillus.* Average nucleotide identity (ANI) calculations with other *Lactobacillus* genomes were above the 95-96% ANI suggested for species demarcation (21), indicating that these MAGs represent strains of established *Lactobacillus* species (Supplement 3). Clostridia EUB1 clustered with *Pseudoramibacter alactolyticus.* The ANI calculations for this MAG with *P. alactolyticus* and related members of the Eubacteriaceae indicated that EUB1 could represent a new genus within the Eubacterium (Supplement 3). The EUB1 MAG is likely the same organism as represented by the *Pseudoramibacter* operational taxonomic unit (OTU) in the 16S rRNA-based identification reported earlier (4). The Clostridia LCO1 did not cluster with a specific genus; ANIs of the LCO1 MAG and related organisms suggest this MAG could represent a novel genus within the Lachnospiraceae (Supplement 3), while the 16S rRNA-based analysis misclassified it as belonging to the *Roseburia* genus (4). The three Actinobacteria MAGs, COR1, COR2, and COR3, clustered within the Coriobacteriaceae family. One of these MAGs (COR2) clustered with *Olsenella umbonata* (22), but the ANI calculation did not support this MAG being a representative of the *Olsenella* genus (Supplement 3). The other two MAGs (COR1 and COR3) formed their own cluster within the Coriobacteriacea, but were sufficiently different in ANI calculations to suggest that each represents new genera within the Coriobacteriaceae. Phylogenetic classification of the other MAGs obtained in this study are provided in Supplement 1.

**Fig. 3.**
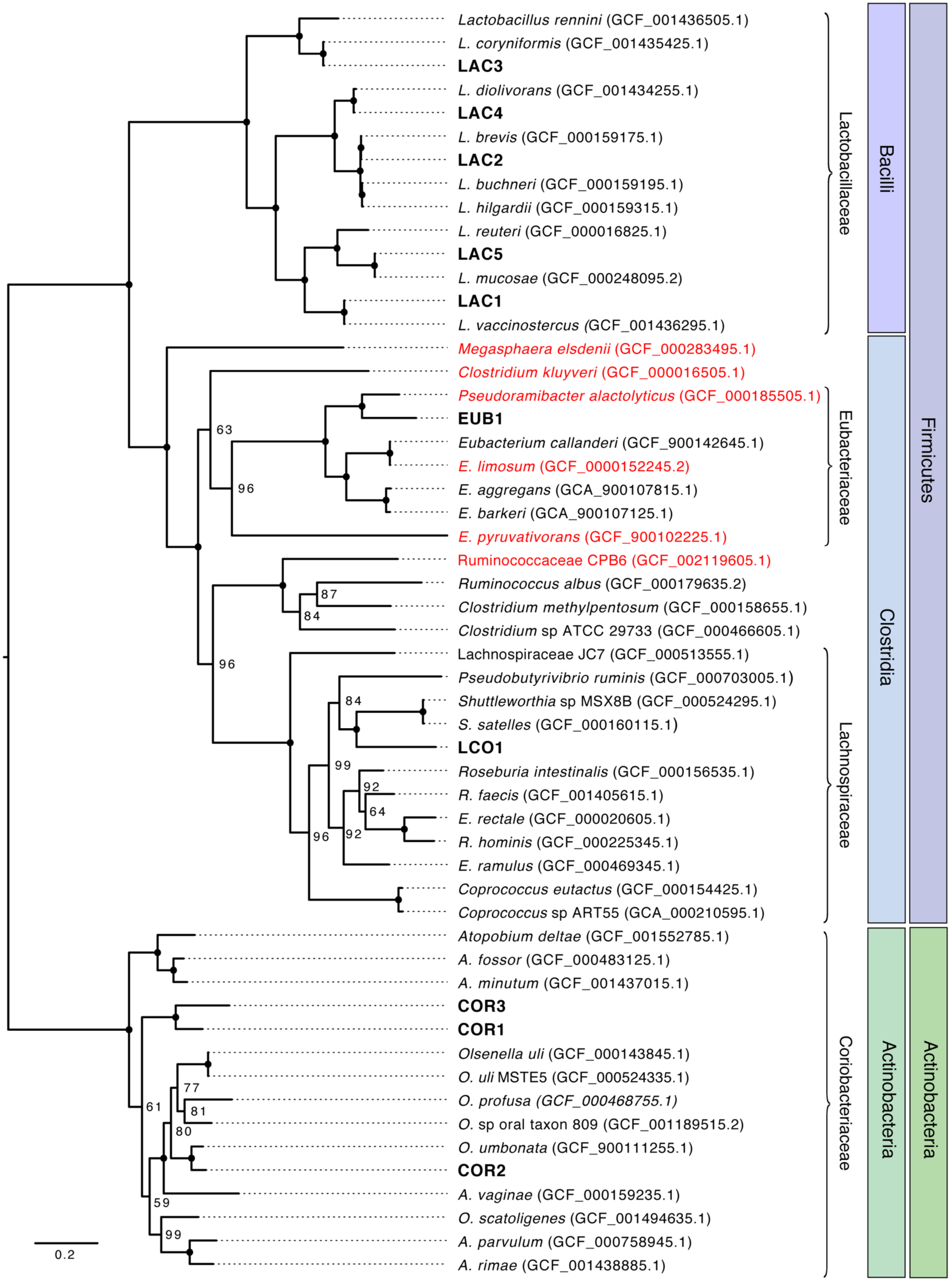
Phylogenetic analysis for ten MAGs obtained from reactor biomass. Draft genomes from this study are shown in bold text. Red text indicates an organism that has been shown to produce MCFA. National Center for Biotechnology Information assembly accession numbers are shown in parentheses. Node labels represent bootstrap support values with solid circles representing a bootstrap support value of 100. The phyla and class of genomes are shown in shaded boxes and families are indicated by brackets For Actinobacteria genomes, Actinobacteria is both the phylum and class.

### Genomic predictions of chemical transformations in the microbiome

A prediction of metabolic networks in the microbiome was performed by analysis of gene annotations or each of the abundant MAGs, whereas expression of the metabolic network was analyzed by mapping mRNA reads to the open reading frames (ORFs) within each of the ten most abundant bacteria. Metabolic reconstruction was performed with automated prediction algorithms (23) and manual curation, particularly of proposed sugar utilization, fermentation and chain elongation pathways (Supplement 4) (1). This analysis identified a set of genes that could be used to model the metabolic potential of the microbiome and also a set of genes with high expression levels in the metatranscriptome. These gene sets were used to analyze the metabolic potential of the microbiome to [1] degrade complex carbohydrates remaining in ethanol conversion residue; [2] transform simple sugars into the fermentation products acetate, lactate, and ethanol; and [3] produce butyrate (C4), C6, and C8 from sugars and fermentation products. The predictions for each of these processes are summarized below.

#### Degradation of complex carbohydrates

Carbohydrates were a large portion of the organic substrates present in the ethanol conversion residue fed to the bioreactor, and uncharacterized carbohydrates. Quantitative analyses indicated that xylose was the most abundant monosaccharide in the residue, accounting for 22% of the organic matter. Glucose was undetected in most samples or a minor component, and other carbohydrates corresponded to 20% of the organic matter in the residue (see CR bar in Fig. 1). Approximately 40% of the uncharacterized carbohydrates were being degraded at the time the metatranscriptomic samples were obtained (Day 96; Fig. 1).

To investigate the expression of genes related to degradation of complex carbohydrates, we analyzed the predicted MAG ORFs using the carbohydrate-active enzyme (CAZyme) database (24). Of particular interest was production of predicted extracellular enzymes that hydrolyze glycosidic bonds in complex carbohydrates, as these may release sugars that can be subsequently metabolized by community members that do not express complex carbohydrate degrading enzymes. The subcellular localization software, CELLO, was used to predict whether individual CAZyme proteins were located within the cytoplasm or targeted to the extracellular space (Supplement 5) (25).

This analysis showed that transcripts encoding genes for several types of glycoside hydrolases (GHs) were abundant in several MAGs in the microbiome (Fig. S1). All LAC MAGs expressed genes encoding extracellular CAZymes that cleave glycosidic bonds between hexose and pentose moieties in xylans. In particular, LAC1 LAC2, and LAC4 expressed genes that encode several extracellular exo-β-xylosidases that could remove terminal xylose molecules from xylans present in the conversion residue (GH43 and GH120; Fig. S1; Supplement 5). LAC2 also had high levels of transcripts for an exo-α-L-1,5-arabinanase (GH93), predicted to release other pentose sugars from arabinan, which accounts for 3% of the sugar polymers in switchgrass (26, 27). In addition, the COR1, COR3 and LAC4 members of the community had high transcript levels for three extracellular CAZymes (GH13) that are predicted to degrade a variety of glucans that may be remaining in switchgrass conversion residue (28). In sum, this analysis predicts that at the time of sampling glucans were degraded by populations represented by *Lactobacillus* and Coriobacteriaceae MAGs, where the populations represented by the LAC MAGs may also have degraded xylans and arabinans. It further suggests that this microbiome is capable of releasing oligosaccharides and sugar monomers from glucans, xylans, and arabinans, the primary components of switchgrass and other plant biomass. The analysis also predicts that LCO1 and EUB1 are not participating in complex carbohydrate degradation.

Bacterial oligosaccharide hydrolysis can also occur in the cytoplasm. All MAGs in this microbiome contained predicted cytoplasmic GH13 enzymes, which are known to degrade hexose oligosaccharides. The microbiome also contained abundant transcripts for genes encoding predicted cytoplasmic CAZYmes that degrade maltose (GH4, GH65), a glucose dimer that may result from extracellular breakdown of glucans (Fig. S1). Transcripts encoding known or predicted cytoplasmic β-glucosidases (GH1, GH3) and β-galactosidases (GH2) are found across the MAGs (Fig. S1). In addition, transcripts that encode β-xylosidases (GH1, GH3) and α-L-arabinofuranosidases (GH2), are found in all the LAC MAGs except LAC3 (Fig. S1). Based on the metatranscriptomic analysis, other cytoplasmic CAZymes predicted to hydrolyze pentose-containing oligosaccharides are predicted to be expressed by the LAC1, LAC2, LAC4, and LAC5 members of this microbiome (Fig. S1).

#### Transport and production of simple fermentation products from sugars

Simple sugars are abundant in ethanol conversion residue and produced during complex carbohydrate hydrolysis. Sugars are therefore expected to be a major substrate for the microbiome. Despite the use of a yeast strain that was engineered for improved xylose utilization in the ethanol fermentation, xylose was the major abundant monosaccharide present in the remaining conversion residue (CR; Fig. 1). As discussed above, the relative transcript levels of genes encoding extracellular GHs (Fig. S1) by several MAGs in the microbiome predict that additional pentoses and hexoses may be released through degradation of complex carbohydrates.

We therefore analyzed the genomic potential of the community to transport sugars, and metabolize them to fermentation products, particularly the known MCFA precursors lactate, acetate, and ethanol. To investigate the ability of the community to transport sugars, MAG ORFs were annotated with the Transporter Classification Database (Supplement 6). Expression of genes associated with the pentose phosphate pathway, phosphoketolase pathways, and glycolysis (Fig. 4a) was analyzed to predict the potential for sugar metabolism within individual MAGs.

**Fig. 4.**
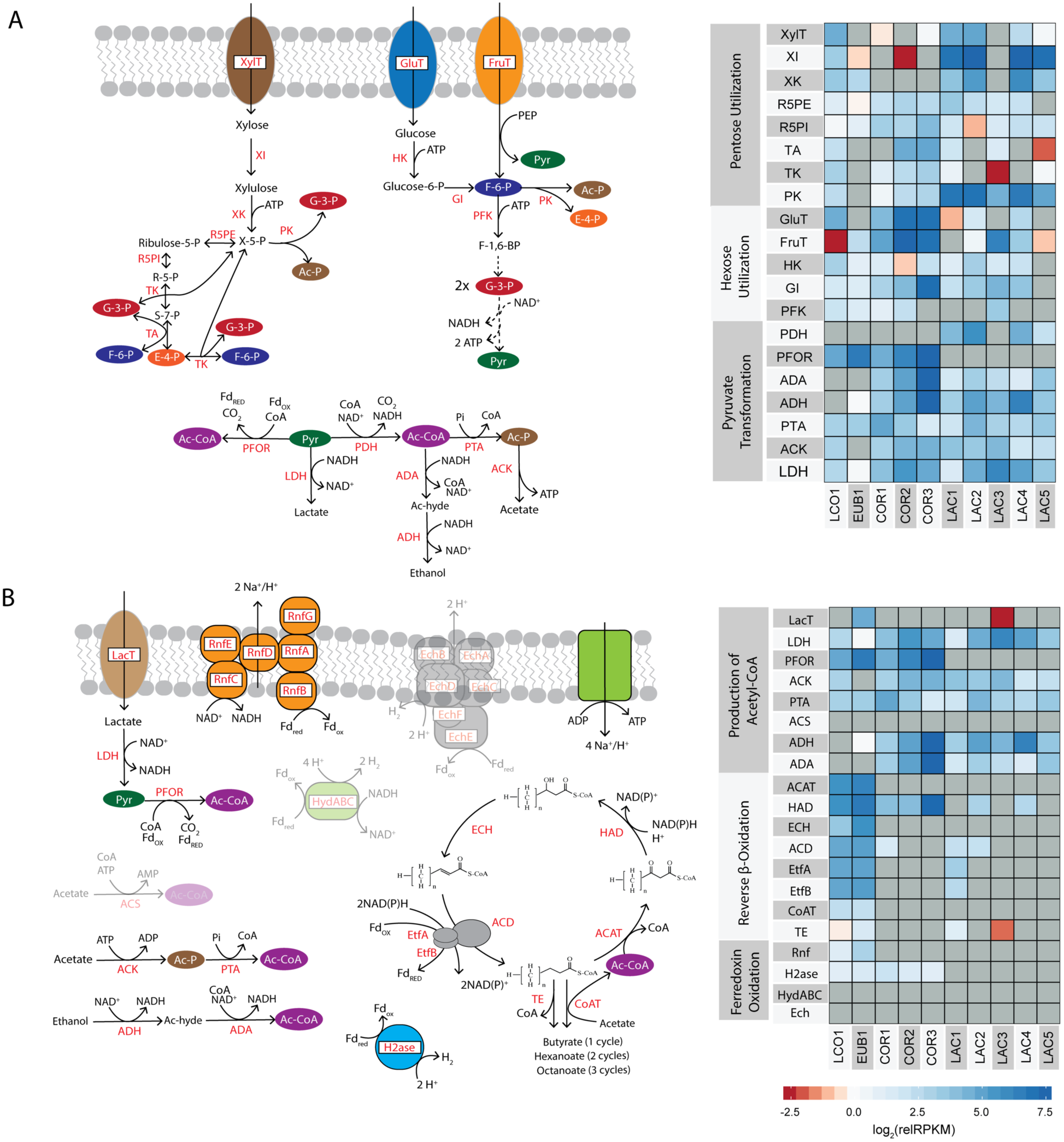
Relative expression of genes involved in the conversion of xylose and lactate to MCFA. Expression of key enzymes involved in (**A**) the utilization of xylose, and **(B)** acyl chain elongation. Dashed lines represent the existence of multiple enzyme reactions between the indicated molecules. The relative RPKM (relRPKM) is normalized to the median RPKM for the MAG. Gene expression data is presented as the log_2_(relRPKM). The color intensities of the heatmaps represent the relative gene expression, with red color intensity indicating expression below median levels and blue intensity indicating expression above median levels. Grey indicates that the gene is absent from the genome. For genes that are not predicted to be expressed by any MAGs, the associated enzyme product is greyed out in the pathway figure. Key pathway intermediates include xylulose, xylulose-5-phosphate (X-5-P), ribose-5-phosphate (R-5-P), sedoheptulose-7-phosphate (S-7-P), erythyrose-4-phosphate (E-4-P), glyceraldehyde-3-phosphate (G-3-P), fructose-6-phosphate (F-6-P), acetate (Ac), acetyl-phosphate (Ac-P), acetyl-CoA (Ac-CoA), lactate and ethanol. Enzyme abbreviations are as follows: **(A)** XylT = xylose transporter, XI = xylose isomerase (EC 5.3.1.5); XK = xylulose kinase (EC 2.7.1.17), R5PE = ribulose-5-phosphate epimerase (EC 5.1.3.1), R5PI = ribose-5-phosphate isomerase (EC 5.3.1.6), TA = transaldolase (EC 2.2.1.2), TK = transketolase (EC 2.2.1.1), PK = D-xylulose 5-phosphate/ D-fructose 6-phosphate phosphoketolase (EC 4.1.2.9), GluT = glucose transporter, FruT = fructose PTS transporter, HK = hexokinase (EC 2.7.1.1, EC 2.7.1.2), G6PI =glucose-6-phoshphate isomerase (EC 5.3.1.9), PFK = phosphofructokinase (EC 2.7.1.11), PDH = pyruvate dehydrogenase complex (EC 1.2.4.1, EC 2.3.1.12, EC 1.8.1.4), PFOR = pyruvate flavodoxin oxidoreductase (EC 1.2.7.-), ADA = acetaldehyde dehydrogenase (EC 1.2.1.10), AD = alcohol dehydrogenase (EC 1.1.1.1), PTA = phosphate acetyltransferase (EC 2.3.1.8), ACK = acetate kinase (EC 2.7.2.1), LDH = lactate dehydrogenase (EC 1.1.1.27), **(B)** LacT = lactate permease, ACS = acetyl-CoA synthetase (EC 6.2.1.1), ACAT = Acetyl-CoA C-acyltransferase (EC 2.3.1.16, EC 2.3.1.9), HAD = 3-Hydroxyacyl-CoA Dehydrogenase (EC 1.1.1.157, 1.1.1.35), ECH = Enoyl-CoA Hydratase (EC 4.2.1.55, EC 4.2.1.17), ACD = Acyl-CoA Dehydrogenase (EC 1.3.99.2, EC 1.3.99.-), EtfA = electron transfer flavoprotein A, EtfB = electron transfer flavoprotein B, TE = thioesterase (EC 3.1.2.20), CoAT = 4-Hydroxybutyrate CoA Transferase (EC 2.8.3.-), Rnf = ferredoxin: NAD^+^ oxidoreductase - Na+ translocating (EC 1.18.1.8), H2ase = ferredoxin hydrogenase (EC 1.12.7.2), HydABC = bifurcating [Fe-Fe] hydorgenase (EC 1.12.1.4), Ech = energy conserving hydrogenase (EchABCDEF)

This analysis found that transcripts from genes encoding predicted carbohydrate transporters were among the most highly abundant mRNAs across the microbiome, accounting for 5.8% of the total transcripts. These putative transporters belonged to a variety of families, including many associated with the ATP-binding cassette (ABC) superfamily, and the phosphotransferase system (PTS) family (TD 4.A.-) (Fig. S2). LCO1, LAC1, LAC2, and LAC3 are predicted to contain xylose transporters (XylT) (Fig. 4a), while glucose (GluT), fructose (FruT), and other hexose transporters were expressed across the LAC, COR, and LCO MAGs (Fig. 4a; Fig. S2). EUB1 only encoded transcripts encoding carbohydrate transporters for uptake of fructose and sucrose (Fig. S2). Overall, this analysis predicts that all MAGs have the potential to transport hexose sugars into the cell, while gene expression patterns observed for the LCO1 and the *Lactobacillus* MAGs (excluding LAC3) predicted that at the time of sampling they played a major role in pentose utilization in this microbiome.

We also analyzed the metatranscriptomic data to investigate potential routes for sugar metabolism. Once transported to the cytoplasm, glucose can be phosphorylated with hexokinase (HK) and converted to fructose-6-phosphate (F-6-P) by glucose-6-phosphate isomerase (GI). Transcripts encoding predicted HK and GI enzymes are abundant for all MAGs within the microbiome (Fig. 4a), except LAC5 for which the assembly does not show homologues of these proteins. Fructose utilization starts with phosphorylation during transport (Fig. 4a). Fructose-6-phosphate (F-6-P) is either phosphorylated to fructose-1,6-bisphosphate (F-1,6-BP) by phosphofructokinase (PFK) in glycolysis or is cleaved to acetyl-P (Ac-P) and erythrose-4-P (E-4-P) by phosphoketolase (PK). While LAC1, LAC2, LAC4, LAC5 and COR3 all lack homologues of genes encoding PFK (a highly-conserved glycolysis enzyme known to be a major target for regulatory control in hexose utilization) (29), they all contain transcripts for homologues of PK (Fig. 4a). In sum, these analyses predict that all of the abundant MAGs in this microbiome can utilize hexoses that may be produced during hydrolysis of complex oligosaccharides.

Transcripts predicted to encode enzymes to convert xylose to xylulose-5-phosphate, xylose isomerase (XI) and xylulose kinase (XK) (30), were abundant in most of the *Lactobacillus* MAGs and LCO1, and either absent or showed very low abundance in LAC3, EUB1 and the COR MAGs (Fig. 4a). Once produced, xylulose-5-P can be degraded through either the phosphoketolase pathway or the pentose phosphate pathway. Transcripts from a gene predicted to encode the diagnostic enzyme of the phosphoketolase pathway, phosphoketolase (PK), which splits xylulose-5-P (X-5-P) into acetyl-P (Ac-P) and glyceraldehyde-3-P (G-3-P), were among the most abundant mRNAs in the *Lactobacillus* MAGs and is also present at high levels in LCO1, accounting for 1.5% of the total transcripts (Fig. 4a; Supplement 4). LCO1 and LAC1 also contained transcripts from homologues of all of the genes needed for the pentose phosphate pathway (R5PE, R5PI, TA, TK in Fig. 4a). Overall, this analysis predicted that multiple routes of pentose utilization could be utilized by the MAGs in this microbiome.

The predicted routes for both hexose and xylose metabolism in this microbiome lead to pyruvate production (Fig. 4a), so we also analyzed how this and other fermentation products might lead to MCFA production in this community. All MAGs contained transcripts encoding lactate dehydrogenase homologues (LDH) (Fig. 4a), an enzyme which reduces pyruvate to lactate. Transcript analysis also predicts that all of the MAGs (except LAC3) can oxidize pyruvate to acetyl-CoA, utilizing either pyruvate dehydrogenase (PDH) or pyruvate flavodoxin oxidoreductase (PFOR) (Fig. 4a). All MAGs (except EUB1) contain transcripts encoding homologues of acetate kinase (ACK), which converts acetyl-phosphate (Ac-P) to acetate while producing ATP (Fig. 4a). Based on predictions of the gene expression data, the COR and LAC MAGs are also able to convert acetyl-CoA (Ac-CoA) to ethanol with aldehyde dehydrogenase (ADA) and alcohol dehydrogenase (ADH). In summary, analysis of the gene expression patterns in the conversion residue microbiome predicts that the MAGs in the LCO, LAC and COR ferment sugars to acetate and lactate, while the LAC and COR members produce ethanol as an additional fermentation product.

#### Elongation of fermentation products to MCFAs

Based on the above findings, we analyzed the microbiome gene expression data to predict which members of the microbiome had the potential for conversion of predicted fermentation products to MCFA. The Clostridia (LCO1 and EUB1) are the only MAGs that contained genes encoding homologues of genes known to catalyze chain elongation reactions in the reverse β-oxidation pathway (Fig. 4b). Thus, the subsequent analysis is based on the prediction that only LCO1 and EUB1 are the major producers of MCFA in this microbiome. Furthermore, based on the analysis of sugar utilization above, we predict that LCO1 is the only microorganism in the community that can directly utilize sugars for MCFA production.

Acetate, lactate and ethanol, are all fermentation products that would require transformation to acetyl-CoA before being used as a substrate for elongation by the reverse β-oxidation pathway. Acetate could be converted to acetyl-CoA utilizing ATP via acetyl-CoA synthase (ACS) or the ACK and phosphate acetyltransferase (PTA) route (Fig. 4b). Alternatively, acetate can be converted to acetyl-CoA with a CoA transferase (CoAT) which transfers a CoA from one carboxylic acid to another (e.g., from butyryl-CoA to acetate, producing butyrate and acetyl-CoA) (Fig. 4b). Genes encoding homologues of ACS and ACK were not found in EUB1, but LCO1 contained abundant transcripts that encoded homologues of both ACK and PTA (Fig. 4a). Both MAGs also contained transcripts predicted to encode CoAT enzymes (Fig. 4b). Taken together, this analysis predicts that acetate may be used as a substrate for MCFA production by LCO1 and EUB1.

Lactate has been proposed as a key intermediate in other microbiomes producing MCFA (12). While transcripts encoding genes for lactate production were abundant in the microbiome (Fig. 4a), lactate did not accumulate to detectable levels during steady operation, but transiently accumulated when the bioreactor received a higher load of conversion residue (4). Transcripts for a gene encoding a predicted lactate transporter (LacT) were abundant in EUB1. In addition, the assembly of LCO1 did not reveal the presence of lactate transporter genes in this MAG, suggesting that only EUB1 may utilize the lactate produced by other MAGs. Neither EUB1 nor LCO1 accumulated transcripts encoding a predicted ADA homologue, which would be required for conversion of acetaldehyde to acetyl-CoA during utilization of ethanol (Fig. 4b). This indicates that if ethanol is produced in this microbiome, it is not used as a significant substrate for MCFA production. Moreover, since ethanol did not accumulate in the reactor during either steady state operation (Fig. 1) or after a high load of conversion residue (4), we predict that ethanol is not a substrate for MCFA production in this microbiome. Rather, based on the predicted activity of LAC and COR MAGs producing lactate and that of EUB1 consuming lactate, we predict that lactate is a key fermentation intermediate for MCFA production.

Within the reverse β-oxidation pathway (Fig. 4b), a key enzyme is an electron-bifurcating acyl-CoA dehydrogenase (ACD) containing two electron transfer flavoproteins (EtfA, EtfB) that pass electrons from NADH to ferredoxin (Fig. 4b) (31). This electron bifurcating complex has been recognized as a key energy conserving mechanism in strictly anaerobic bacteria and archaea (17, 31) and studied in detail in butyrate producing anaerobes (32, 33). Transcripts for genes encoding a homologue of the acyl-CoA dehydrogenase complex (ACD, EtfA, EtfB) were abundant in both LCO1 and EUB1, as are those from other genes predicted to be involved in this pathway (Fig. 4b). Chain elongation by the reverse β-oxidation pathway conserves energy by increasing the ratio of reduced ferredoxin (a highly electropositive electron carrier) to the less electropositive NADH (1). In organisms that use this pathway, oxidation of ferredoxin by the Rnf complex generates an ion motive force, and ATP synthase utilizes the ion motive force to produce ATP (17). We found that transcripts for genes encoding homologues of all six subunits of the Rnf complex were abundant in both EUB1 and LCO1 (RnfABCDEG, Fig. 4b). To maintain cytoplasmic redox balance, reduced ferredoxin could transfer electrons to H^+^ via hydrogenase, generating H_2_. LCO1 and EUB1, along with the COR MAGs, contained abundant transcripts for genes predicted to produce ferredoxin hydrogenase (H2ase, Fig. 4b), supporting the hypothesis that H_2_ production plays a role within this MCFA-producing microbiome. We also looked for two additional hydrogenases known to conserve energy either through the translocation of protons (EchABCDEF Fig. 4b,) or by electron confurcation, utilizing electrons from both NADH and reduced ferredoxin (HydABC, Fig. 4b) (17). It does not appear that these systems play a major role in H_2_ production in this microbiome since none of the MAGs contained genes encoding homologues of the known components for either of these enzyme complexes (Fig 5b).

### Thermodynamic analysis of MCFA production in the microbiome

The above analysis predicted several potential routes for MCFA production by LCO1 and EUB1 in this microbiome. To evaluate the implications of these potential chain elongation routes, we used thermodynamic analysis to investigate the energetics of the predicted transformations. For this, we reconstructed metabolic pathways for xylose and lactate conversion, as well as ATP yields based on the data obtained from gene expression analyses (Tables S2-S3, Supplement 7). Metabolic reconstructions considered xylose (Table S2) and lactate (Table S3) as major substrates for synthesis of C4, C6, and C8 products. In addition, both LCO1 and EUB1 have the potential to use a CoAT or a thioesterase (TE) as the terminal enzyme of the reverse β-oxidation pathway (Fig. 4b), so we considered both possibilities in the thermodynamic analysis. We used these reconstructions to calculate the free energy changes of the overall biochemical reactions assuming an intracellular pH of 7.0, a temperature of 35°C, and H_2_ partial pressures of 1.0 x 10^−6^, 1.0, and 6.8 atm for low, standard, and high H_2_ partial pressure, respectively. The low value is the approximate concentration of H_2_ in water that is in equilibrium with the atmosphere and the high value is an expected maximum in a pressurized mixed culture fermentation system (34). We also compared the efficiency of ATP production to an expected maximum yield of 1 ATP per −60 kJ energy generated by the overall chemical transformation (35).

The use of xylose as the substrate (Table S2, Eqs. 3-8) is possible for LCO1 but not EUB1, since the later MAG lacks genes to transport and activate xylose to xylulose-5-P (XylT, XI, XK, Fig. 4a). Our analysis predicts that, with a pathway containing a terminal CoAT enzyme, the ATP yield (mol ATP mol^−1^ xylose) does not increase if longer chain MCFAs are produced. However, if TE is used for the terminal step of reverse β-oxidation (Eqs. 6-8), the overall ATP yield is lower, but it increases with increasing product length, and C8 production provides a 17% increase in ATP yield versus production of C4. This suggests that LCO1 has no energetic benefit for producing C6 or C8 solely from xylose unless TE is used as the terminal enzyme of reverse β-oxidation. Additionally, the higher ATP yield of xylose conversion to C4 (Eqs. 3 and 6), in comparison to xylose conversion to lactate and acetate by other members of the microbiome (Eq. 2), may explain why LCO1 reached higher abundance in the microbiome compared to the other less abundant MAGs (LAC) that are predicted to ferment xylose to lactate and acetate (Fig. 2). In production of C4 and C8, no H_2_ is predicted to be formed if a CoAT is utilized (Eqs. 3 and 5), whereas H_2_ production is predicted when C6 is produced (Eq. 4). On the other hand, if a TE terminal enzyme is utilized for the reverse β-oxidation, H_2_ is predicted to be produced for all carboxylic acid products.

Additional metabolic reconstructions analyzed the co-utilization of xylose with a monocarboxylic acid (Table S2, Eq 9-18). This analysis predicted that co-metabolism of these substrates could provide an energetic advantage (i.e., higher mol ATP per mol of xylose) if H_2_ is utilized as an electron donor. This suggests that H_2_, produced by either EUB1 or COR MAGs (H2ase, Fig5a), may be utilized by LCO1 to support MCFA production. If TE is used as the terminal enzyme of reverse β-oxidation (Eqs. 14-18), there is no increase in ATP yield versus utilization of xylose as the sole carbon source (Eqs. 6-8).

We also modeled MCFA production from lactate by EUB1, since the gene expression data suggested that EUB1 could transform lactate to MCFA. In models utilizing CoAT (Table S3, Eq. 19-21) as a final step in MCFA production, the ATP yield increases as longer chain MCFA are produced, but the free energy released is near the expected limit for ATP production (35) under conditions of low H_2_ partial pressure and below this limit at high H_2_ partial pressures (Table S3). If TE is utilized as a final step in MCFA production by EUB1 (Eq. 22-24), lower ATP yields are predicted, and in that case the production of longer chain MCFAs has a more pronounced effect on the ATP generated per mol of lactate consumed.

For instance, production of C6 results in a 100% increase in the ATP yield compared to producing C4. However, each elongation step reduces the amount of energy released per mol ATP produced, such that production of C8 from lactate results in the release of −58 kJ per ATP produced under high H_2_ conditions, which is near the expected limits for a cell to conserve chemical energy as ATP. Overall, the thermodynamic analysis does not unequivocally predict which terminal enzyme may be energetically more advantageous for MCFA production from lactate. While using TE would result in more favorable free energy release than when using CoAT, the predicted ATP yields are lower with TE than with CoAT. We also note that although CoAT transcript abundance was higher than TE transcript abundance (Fig. 4b), expression alone cannot be used as a predictor of which terminal enzyme was primarily used since a kinetic characterization of these enzymes is not available. Regardless, the thermodynamic modeling predicts that, in all conditions, H_2_ will be produced during lactate elongation (Table S3), and that TE could be a better terminal enzyme to force production of longer chain acids in order to maximize ATP yield (Table S3).

When modeling scenarios utilizing lactate plus carboxylic acids as growth substrates (Table S3, Eqs. 25-36), their elongation by EUB1 would increase the amount of ATP it could produce compared to using lactate only if using a terminal CoAT (Eqs. 19-21). H_2_ production or consumption is not predicted in these scenarios, and the calculated free energy released per mol of ATP produced (−50 to −53 kJ mol^−1^ ATP) is low, near the physiological limit of −60 kJ mol^−1^ ATP for energy conservation by the cell. Models with TE as the terminal enzyme in reverse β-oxidation were also analyzed (Eqs. 31-36) even though EUB1 is not predicted to have this ability as it lacks ACS and ACK needed to utilize acetate (Fig. 4b). In such models, producing C6 and C8 from lactate plus acetate (Eqs. 32-33) is energetically favorable, whereas C4 production is not (Eq. 31).

## Discussion

In this study, we illustrate how combining genomic, computational and thermodynamic predictions can illustrate how a microbial community can convert organic substrates in lignocellulosic conversion residues into MCFA (Fig. 5). Specifically, this approach predicts that the coordinated and step-wise metabolic activity of different members of this microbiome allow cleavage of complex five- and six-carbon containing polysaccharides; conversion of sugars into simple fermentation products; and utilization of sugars and intermediate fermentation products for MFCA production. This approach further predicts the role of intracellular and extracellular reductants in these processes. Below, we illustrate the new insight that has been gained on the activity of a MCFA producing microbiome and how this might relate to other systems.

**Fig. 5.**
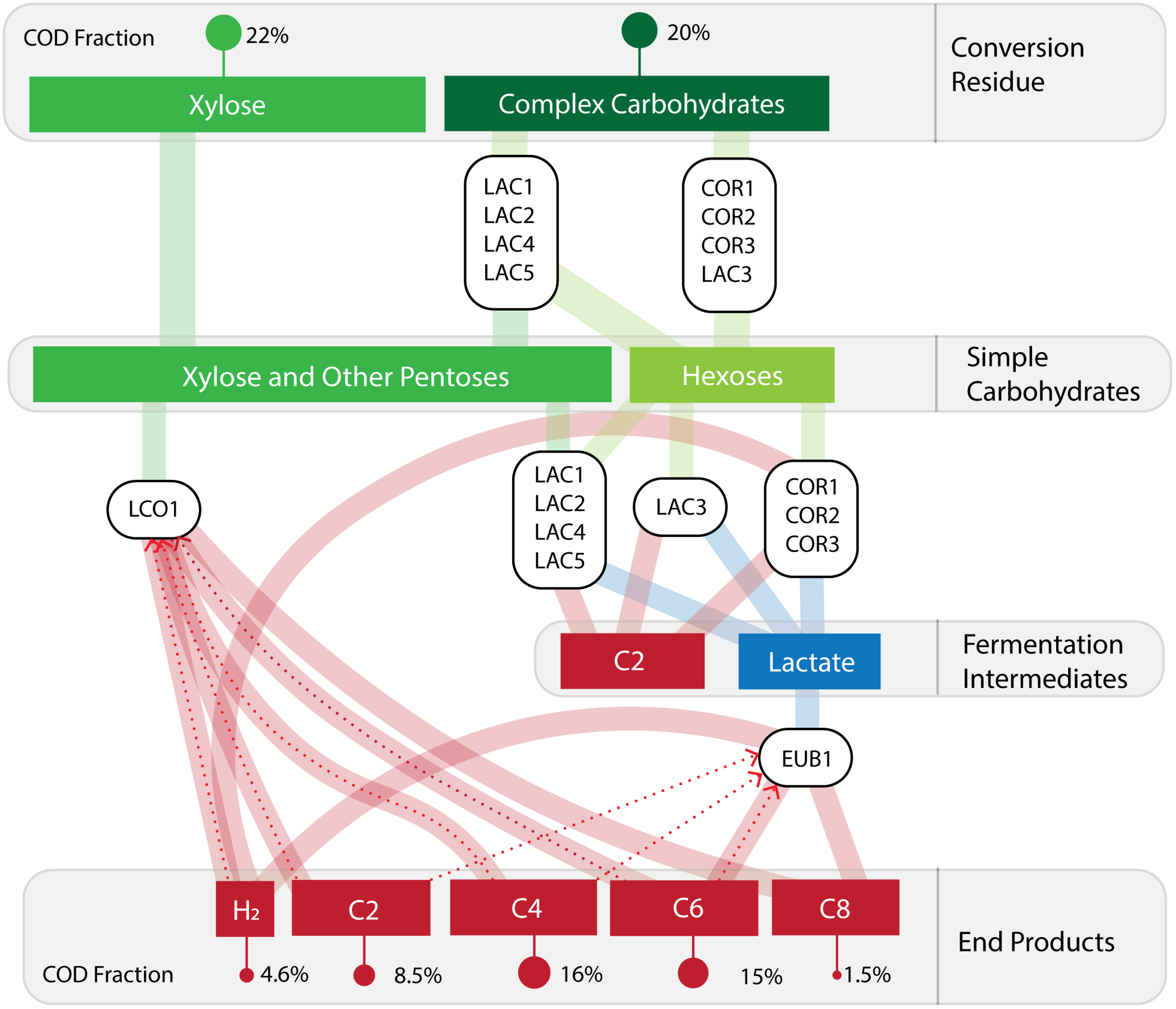
Predicted transformations of major substrates in conversion residue to MCFA by this anaerobic microbiome. The microbes in the LAC and COR bins are predicted to produce sugars from complex carbohydrates. Simple carbohydrates, including xylose remaining in conversion residue, are converted to lactate and acetate by *Lactobacillus* (LAC) and Coriobacteriaceae (COR) MAGs. The Lachnospiraceae (LCO1) MAG converts pentoses directly to butyric acid (C4). The Eubacteriaceae (EUB1) produces hexanoic acid (C6) and octanoic acid (C8) from lactate. Further, LCO1 may utilize hydrogen to elongate C2 and C4 to MCFA, as represented by dashed lines. Additionally, EUB1 may elongate C2, C4 and C6 to C8.

The microbial community studied here is similar in phylogenetic composition to other microbial communities producing MCFA. For instance, in a community fed with dilute ethanol and stillage, *Lactobacillus* and a member of the Clostridium Group IV were abundant (12). In a system producing MCFA using thin stillage produced from corn ethanol, *Lactobacillus* were enriched alongside *Megasphaera*, a known MCFA-producing Firmicute (10). In a reactor converting wastes from wine production to MCFA, *Lactobacillus* and Clostridia related to *Ruminococcus* were enriched (9). In each case, a community containing carbohydrate fermenting organisms and potential MCFA producing organisms emerged.

Our data suggest that the contribution of *Lactobacillus* in this microbiome is in extracellular carbohydrate degradation and subsequent metabolism of pentose-and hexose-containing carbohydrates, while Coriobacteriaceae are predicted to metabolize hexose-containing carbohydrates. The combined metabolic activities of these two MAGs would produce oligosaccharides and monomeric sugars that would become available to these and other members of the microbiome. Metabolic reconstruction combined with microbiome transcript levels also suggest that the *Lactobacillus* and Coriobacteriaceae MAGs produce fermentation end products, primarily lactate and acetate, from these carbohydrates. Coriobacteriaceae, however, are also predicted to produce H_2_. In addition, microbiome gene expression patterns indicate that two MAGs, EUB1 and LCO1, produce MCFA via reverse β-oxidation. LCO1 is predicted to consume xylose based on gene expression analysis, whereas RNA abundance measurements indicate that EUB1 consumes lactate.

We used thermodynamics to analyze hypothetical scenarios of MCFA production by EUB1 and LCO1. Although the comparison of these hypothetical scenarios did not provide an unequivocal answer to how chain elongation occurs in LCO1 and EUB1, it is helpful to generate hypotheses that could eventually be tested in future research. Our thermodynamic analysis predicts that the most energetically advantageous metabolism for LCO1 (based on ATP production per mol xylose consumed) is the consumption of xylose, H_2_ and carboxylates to produce C4, C6, and C8 while utilizing CoAT as a terminal enzyme. While xylose is a major component of conversion residue (CR, Fig. 1), H_2_ is expected to be produced by Coriobacteriaceae MAGs and EUB1. For EUB1, which is expected to utilize lactate, our analysis predicts that production of MCFA produces higher amounts of ATP, with C6 resulting in a 2-fold increase in ATP production versus producing C4 when consuming lactate as a sole substrate.

Predictions from our thermodynamic modeling indicate that C8 production from lactate is energetically advantageous. However, this is at odds with C6 being produced from conversion residue at higher concentrations than C8 (Fig. 1). It is known that C8 is a biocide, so it may be that C8 accumulation is limited by the level of tolerance community members have for this product (12). It is also possible that higher C6 production indicates a more important role of C6 production by LCO1 without lactate being an intermediate metabolite. It has also been shown that removal of C8 allows for higher productivities of carboxylate platform systems (1).

H_2_ production and interspecies H_2_ transfer are known to have significant impacts on the metabolism of microbial communities (36). Our analysis predicts a role of H_2_ in supporting chain elongation in a carboxylate platform microbiome. While high H_2_ partial pressures are proposed to inhibit production of acetate and other carboxylic acids (37, 38), organisms that use the phosphoketolase pathway (the *Lactobacillus* and LCO1 MAGs identified in this study) can produce C2, C4 and C8 without producing H_2_ (Table S2, Eqs. 2, 3 and 5). While conversion of lactate to MCFA (Table S3, Eqs. 19-24) is predicted to produce H_2_, other processes such as co-utilization of xylose and a monocarboxylic acid for MCFA production (Table S2, Eq. 9-18) would consume H_2_. Therefore, H_2_ accumulation is not expected to limit production of MCFA, although H_2_ partial pressures may influence the metabolic routes utilized by the microbiome.

In considering how to further improve the production of MCFA with a microbiome, additional work is needed to characterize and engineer reverse β-oxidation proteins from the Firmicutes in order to improve production of organic acids longer than C4. Further, our data predict that the terminal enzyme of reverse β-oxidation can influence production of MCFA. While a CoAT enzyme results in higher ATP production, a TE makes production of MCFA more energetically advantageous by increasing the ATP yield for production of C6 and C8 compared to C4 (Tables S2-S3). Therefore, engineering chain-elongating organisms to only have a TE rather than CoAT may improve production of MCFA.

Our metabolic reconstructions predict that lactate was a key fermentation product that supports MCFA production. Therefore, strategies to enhance lactate production and minimize other fermentation products (fermentation of carbohydrates to lactate, rather than acetate in this example), could improve production of desired end products. Moreover, designing strategies to enrich a community that produces a critical intermediate like lactate by one pathway (e.g., homofermentative lactate-producing *Lactobacilli*, rather than hetero-fermenters producing both lactate and C2) could improve performance of the microbiome. However, the principles controlling the presence or dominance of heterofermentative versus homofermentative organisms in microbial communities remain largely unexplored. Alternatively, higher production of a desired product, C8, could be achieved by adjusting the abundance or establishing a defined co-culture containing a lactic acid bacterium capable of complex carbohydrate degradation, such as LAC1, and a lactate-elongating organism, such as EUB1. The ability to establish defined synthetic communities, to adjust the abundance of microbiome members or to regulate the metabolic routes within the microbiome may allow more control over the function of a microbiome for production of MCFA or to optimize other traits.

In summary, this work demonstrates that one can dissect and model the composition of microbiomes as a way to understand the contribution of different community members to its function. In the case of an anaerobic carboxylate platform microbiome fed lignocellulosic ethanol conversion residue, two Clostridia-related organisms (EUB1 and LCO1) are predicted to be responsible for production of MCFA via reverse β-oxidation. This provides a genome-centric rationale for the previously established correlation between Clostridia-related abundance and MCFA production noted in carboxylate platform systems (4, 12). This study further predicts that the terminal enzyme in product synthesis and the fermentation end products produced by other community members can play a role in determining predominant products of this microbiome. These approaches, concepts and insights should be useful in predicting and controlling MCFA production by reactor microbiomes and in analyzing the metabolic, genomic and thermodynamic factors influencing the function of other microbiomes of health, environmental, agronomic or biotechnological importance.

## Methods

### Production of conversion residue

Switchgrass used to generate conversion residue was treated by ammonia fiber expansion and enzymatically treated with Cellic CTec3^®^ and Cellic HTec3^®^ (Novozymes) to digest cellulose and hemicellulose (to glucose and xylose, primarily) (39). Hydrolysate was fermented with *Saccharomyces cerevisiae* Y128, a strain with improved xylose utilization (40). Fermentation media was distilled to remove ethanol (4).

### Bioreactor operation

The bioreactor was seeded with acid digester sludge from the Nine Springs Wastewater treatment plant in Madison, WI. The retention time of the semi-continuous reactor was maintained at six days by pumping conversion residue into the reactor, pumping reactor effluent from the reactor once per hour, and maintaining a liquid volume of 150mL in the reactor. The reactor was mixed with a magnetic stir bar. The temperature of the reactor was controlled at 35 °C using a water bath, and the pH of the reactor was maintained at 5.5 by feeding 5M KOH through a pump connected to a pH controller. This reactor sustained MCFA production for 252 days (4).

### Metabolite analysis

Samples from the bioreactor and conversion residue were collected for metabolite analysis. All samples were filtered using 0.22 μm syringe filters (ThermoFisher Scientific SLGP033RS, Waltham, MA, USA). Chemical oxygen demand (COD) analysis was performed using High Range COD Digestion Vials (Hach 2125915, Loveland, CO, USA) per standard methods (41). Soluble carbohydrates were measured with the anthrone method (42). Glucose, xylose, acetic acid, formic acid, lactic acid, succinic acid, pyruvic acid, glycerol and xylitol were analyzed with high performance liquid chromatography and quantified with an Agilent 1260 Infinity refractive index detector (Agilent Technologies, Inc. Palo Alto, CA) using a 300 x 7.8 mm Bio-Rad Aminex HPX-87H column with Cation-H guard (BioRad, Inc., Hercules, CA). Acetamide, ethanol, n-propionic acid, n-butyric acid, iso-butyric acid, n-pentanoic acid, iso-pentanoic acid, n-hexanoic acid, iso-hexanoic acid, n-heptanoic acid, and n-octanoic acid were analyzed with tandem gas chromatography-mass spectrometry. An Agilent 7890A GC system (Agilent Technologies, Inc. Palo Alto, CA) with a 0.25 mm Restek Stabilwax DA 30 column (Restek 11008, Belefonte, PA) was used. The GC-MS system was equipped with a Gerstel MPS2 (Gerstel, Inc. Baltimore, MD) auto sampler and a solid-phase micro-extraction gray hub fiber assembly (Supelco, Bellefonte, PA). The MS detector was a Pegasus 4D TOF-MS (Leco Corp., Saint Joseph, MI). Stable isotope labeled internal standards were used for each of the analytes measured with GC-MS.

### DNA and RNA sequencing

Biomass samples, consisting of centrifuged and decanted 2 mL aliquots, were collected at Day 12, Day 48, Day 84, Day 96, and Day 120 of reactor operations from initial start-up. Samples were also taken at 96 days and flash-frozen in liquid nitrogen for RNA extraction. For DNA extraction, cells were lysed by incubating in a lysis solution (1.5M sodium chloride, 100mM trisaminomethane, 100mM ethylenediamine (EDTA), 75mM sodium phosphate, 1% cetyltrimethylammonium bromide, and 2% sodium dodecyl sulfate(SDS)), lysozyme (Thermo Fisher Scientific, MA, USA), and proteinase K (New England Biolabs,MA, USA). We then added 500 uL of a 24:24:1 solution of phenol: chloroform: isoamyl alcohol and bead-beat samples for 2 minutes. After bead-beating, biomass was centrifuged at 5,000 rcf for 3 minutes and the entire supernatant was transferred to a 1.5 mL centrifuge tube. Samples were centrifuged again at 12,000 rcf for 10 minutes and the aqueous layer was then removed to a new centrifuge tube. A second phase separation was then performed using chloroform. After centrifuging again and separating the aqueous phase, 500 uL of isopropanol was added to each samples and samples were then incubated at −20 deg C for 24 hours. Following this incubation, samples were centrifuged at 12,000 rcf for 30 minutes at 4 deg C, decanted, and washed with 70% ethanol. After air-drying the samples, pellets were resuspended in 100 uL of TE buffer and 2 uL of 10 mg/mL RNAse was added to each sample. Samples were incubated for 15 minutes at 37 deg C. We then added 100uL of a 24:24:1 solution of phenol: chloroform: isoamyl alcohol to each sample and centrifuged at 12,000 rcf for 10 minutes. We separated the aqueous phase to a new centrifuge tube and added 100 Ul of chloroform. Again, samples were centrifuged at 12,000 rcf for 10 minutes and the aqueous phase was separated to a new centrifuge tube. We then added 10 uL of 3M sodium acetate and 250 uL of 95% ethanol to each sample and incubated for 24 hours at −20 deg C. Samples were centrifuged at 12,000 rcf for 30 minutes at 4 deg C and the pellets were washed with 70% ethanol. After air-drying, pellets were resuspended in 50uL of TE buffer. After re-suspending the DNA, quantity, purity, and quality were assessed with a Qubit 4 Fluorometer (Thermo Fisher Scientific, MA, USA), a Nanodrop 2000 spectrophotometer (Thermo Fisher Scientific, MA, USA), and gel electrophoresis.

For RNA extraction, cells were lysed by incubating in a lysis solution (20 mM sodium acetate, 1 mM EDTA, and 0.5% SDS prepared in diethylpyrocarbonate-treated water (Invitrogen, CA, USA)) and TRIzol (Invitrogen, CA, USA). The treated cells were subjected to 2 minutes of bead beating using Lysing Matrix A (MP Biomedicals, CA, USA). After this step, successive phase separations with phenol: chloroform: isoamyl alcohol and chloroform were used to separate nucleic acids from additional cell material, as described above. RNA was further purified with an RNEasy Mini Kit (Qiagen, Hilden, Germany) and on-column DNAse 1 (Qiagen, Hilden, Germany) treatment. After re-suspending the RNA, quantity, purity, and quality were assessed with a Qubit 4 Fluorometer (Thermo Fisher Scientific, MA, USA), a Nanodrop 2000 spectrophotometer (Thermo Fisher Scientific, MA, USA), and gel electrophoresis. RNA samples were submitted to the University of Wisconsin Gene Expression Center for quality control with a Bioanalyzer (Agilent, CA, USA), ribosomal RNA reduction with a RiboZero-Bacteria rRNA Removal Kit (Illumina, CA, USA) with a 1μg RNA input. Strand-specific cDNA libraries were prepared with a TruSeq RNA Library Prep Kit (Illumina, CA, USA).

DNA and RNA were sequenced with the Illumina HiSeq 2500 platform (Illumina, CA, USA). For DNA, an average insert size of 550 bp was used and 2 x 250 bp reads were generated. For RNA, 1 x 100 bp reads were generated. Raw DNA and cDNA reads can be found on the National Center for Biotechnology Information (NCBI) website under BioProject PRJNA418244.

### Metagenomic assembly, binning, and quality control

DNA sequencing reads were filtered with Sickle using a minimum quality score of 20 and a minimum sequence length of 100 (43). Reads from all five samples were then co-assembled using metaspades and kmer values of 21, 33, 55, 77, 99, and 127 (44). Binning of assembled contigs was performed with MaxBin v2.2.1 (45). The quality, completeness, and contamination of each bin was analyzed with CheckM v1.0.3 (46). Read mapping was performed with BBMAP v35.92 (https://sourceforge.net/projects/bbmap) to estimate the relative abundance of each bin. Relative abundance was calculated by normalizing the number of mapped reads to the genome size.

### Phylogenetic analysis

Phylogeny of the draft genomes was assessed using 37 universal single-copy marker genes with Phylosift v1.0.1 (47). In addition to the draft genomes, 62 publically-available genomes of related organisms were used to construct a phylogenetic tree. Concatenated amino acid sequences of the marker genes were aligned with Phylosift, and a maximum likelihood phylogenetic tree was constructed with RAxML v8.2.4 with the PROTGAMMAAUTO model and 100 bootstraps (48). ANI calculations were performed using JSpecies (49).

### Genome annotations

Draft genomes were annotated with MetaPathways v2.5.(23) Open reading frames (ORFs) were predicted using Prodigal v2.0 (50), and the ORFs were annotated with the following databases: SEED (accessed March 2013), Clusters of Orthologous Groups (COG, accessed December 2013), RefSeq (accessed January 2017), Metacyc (accessed October 2011), and KEGG (accessed January 2017). The LAST algorithim was used for assigning functional annotations (51). Functional annotations for each MAG are provided in Supplement 4. Draft genomes were further annotated with the CAZY database (24). CELLO was used to determine the subcellular location of the CAZYs (25). Transporters were identified using the Transporter Classification Database.

### Transcript analysis

Analysis of transcript data was performed as described in Lawson, et al. (52). cDNA reads were quality filtered as described above for DNA. SortMeRNA was used to remove rRNA sequences using multiple databases for RNA sequences (53). The remaining non-rRNA sequences were then mapped back to the draft genomes using BBMap v35.92 (https://sourceforge.net/projects/bbmap) with the minimum sequence identity set to 0.95. Ambiguous reads with multiple top-hit mapping locations were assigned to a random ORF. The number of RNA reads mapping to each ORF was calculated with htseq-count v0.6.1 with the “intersection-strict” parameter (54). Relative gene expression (RPKM) was calculated for each ORF by normalizing the number of mapped RNA reads for each ORF to the ORF length and the total number of RNA reads mapping back to the genome. The relative RPKM (relPKM) was then calculated as the ratio of the RPKM for the ORF to the median RPKM across the draft genome. Finally, the log_2_(relRPKM) was calculated to determine the log-fold difference. As such, a positive number corresponds to greater than median expression levels and a negative number to expression below median levels.

## Acknowledgements

This work was funded by the DOE Great Lakes Bioenergy Research Center (DOE BER Office of Science DE-FC02-07ER64494 and DE-SC0018409). M.J.S. is supported by the National Science Foundation Graduate Research Fellowship Program under grant No. DGE-1256259. C.E.L is supported by a Postgraduate Scholarship–Doctoral (PGS-D) award from the National Sciences and Engineering Research Council of Canada (NSERC) and a Wisconsin Distinguished Graduate Fellowship. JJH was supported by the National Institute of Food and Agriculture, U.S. Department of Agriculture, under award number 2016-67012-24709. The authors thank Francisco Moya for assistance with RNA extraction and the University of Wisconsin Biotechnology Center DNA Sequencing Facility and Gene Expression Center for library preparation and Illumina sequencing services.

---

**Table S1.** Summary of MAGs obtained from DNA sequence analysis of the reactor microbiome. Taxonomy, completeness, and contamination were estimated with CheckM. Draft genomes were assembled from five independent reactor samples. These MAGs represent the ten most abundant MAGs at Day 96 (Fig. 1).

| Bin ID | Taxonomy | Completeness (%) | Contamination (%) | Genome size (bp) | # Scaffolds | N50 | GC (%) | Predicted Genes |
| --- | --- | --- | --- | --- | --- | --- | --- | --- |
| LCO1 | Firmicutes; Clostridia; Clostridiales; Lachnospiraceae; Shuttleworthia | 95.4 | 0.0 | 2,106,912 | 44 | 103,964 | 45.7 | 1,900 |
| EUB1 | Firmicutes; Clostridia; Clostridiales; Eubacteriaceae; Pseudoramibacter | 97.8 | 0.2 | 2,002,609 | 35 | 142,846 | 51.2 | 1,857 |
| COR1 | Actinobacteria; Actinobacteria; Coriobacteriales; Coriobacteriaceae; Olsenella | 99.2 | 0.8 | 2,512,349 | 225 | 22,880 | 59.0 | 2,358 |
| COR2 | Actinobacteria; Actinobacteria; Coriobacteriales; Coriobacteriaceae; Olsenella | 100 | 1.6 | 2,422,853 | 155 | 34,678 | 64.8 | 2,185 |
| COR3 | Actinobacteria; Actinobacteria; Coriobacteriales; Coriobacteriaceae; Olsenella | 98.4 | 7.4 | 3,647,413 | 533 | 13,368 | 55.5 | 4,068 |
| LAC1 | Firmicutes; Bacilli; Lactobacillales; Lactobacillaceae; Lactobacillus | 99.5 | 1.1 | 2,633,889 | 18 | 640,122 | 43.6 | 2,567 |
| LAC2 | Firmicutes; Bacilli; Lactobacillales; Lactobacillaceae; Lactobacillus | 99.4 | 1.6 | 3,179,174 | 79 | 122,889 | 40.5 | 2,989 |
| LAC3 | Firmicutes; Bacilli; Lactobacillales; Lactobacillaceae; Lactobacillus | 99.2 | 1.4 | 2,704,063 | 174 | 29,509 | 43.0 | 2,731 |
| LAC4 | Firmicutes; Bacilli; Lactobacillales; Lactobacillaceae; Lactobacillus | 98.9 | 1.3 | 3,335,227 | 95 | 86,779 | 40.2 | 3,150 |
| LAC5 | Firmicutes; Bacilli; Lactobacillales; Lactobacillaceae; Lactobacillus | 80.1 | 0.8 | 1,487,044 | 181 | 12,363 | 46.1 | 1,524 |

**Fig. S1.**
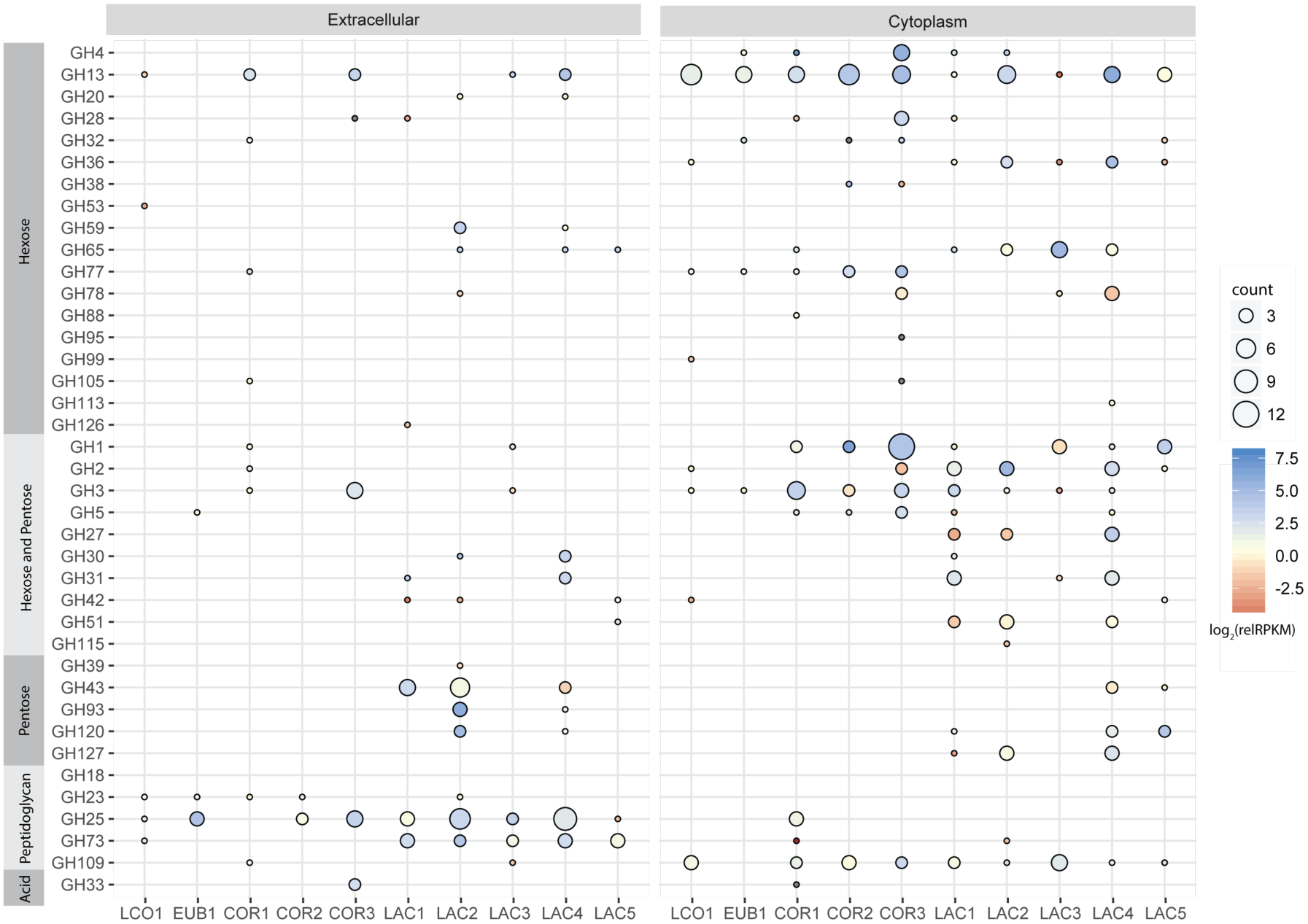
Relative expression of extracellular and cytoplasmic glycoside hydrolase enzymes. Circle size indicates the number of ORFs annotated within each bin as the indicated CAZyme family. CAZyme families are grouped according to the target substrate moieties, being either bonds between hexoses only, bonds between both hexoses and pentoses, bonds between pentoses only, peptidoglycan-specific moieties, or acid moieties. The relative RPKM (relRPKM) is normalized to the median RPKM for the MAG. Gene expression data is presented as the log_2_(relRPKM). The color intensity represents the relative gene expression, with red color intensity indicating expression below median levels and blue intensity indicating expression above median levels. The represented expression is the highest relative expression for an individual gene within the indicated family. Descriptions of each family are available through the CAZY database (http://www.cazy.org/Glycoside-Hydrolases.html).

**Fig. S2.**
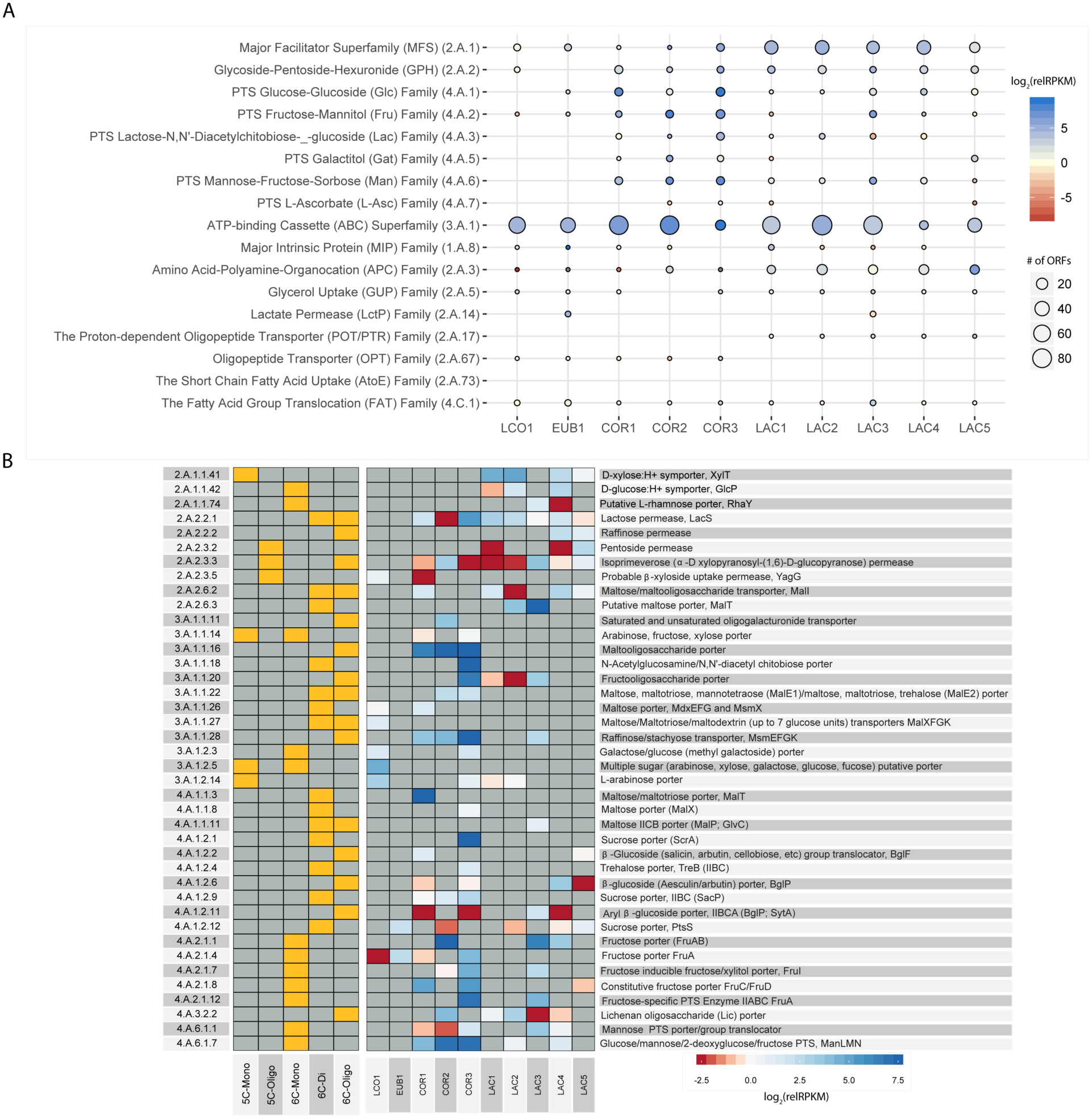
Expression of known or predicted transporters by the 10 most abundant microbiome members. Transporters were annotated with the Transporter Classification Database. **(A)** The number of genes and relative expression for each analyzed MAG are shown for Transporter Database families. Circle size indicates the number of ORFs annotated within each bin as the indicated CAZY class or family. The color intensity represents the relative gene expression according to the shown scale. The represented expression is the highest relative expression for an individual gene within the indicated transporter family. (B) The expected substrates, either a pentose (5C-Mono), pentose-containing oligosaccharide (5C-Oligo), hexose (6C-mono), hexose disaccharide (6C-Di), or hexose-containing oligosaccharide (6C-Oligo) are indicated by orange squares. The color intensity represents the relative gene expression, with red color intensity indicating expression below median levels and blue intensity indicating expression above median levels. The transporter family ID and descriptions are shown. More detailed descriptions of transporters are available from the Transporter Database (http://www.tcdb.org).

**Table S2.**
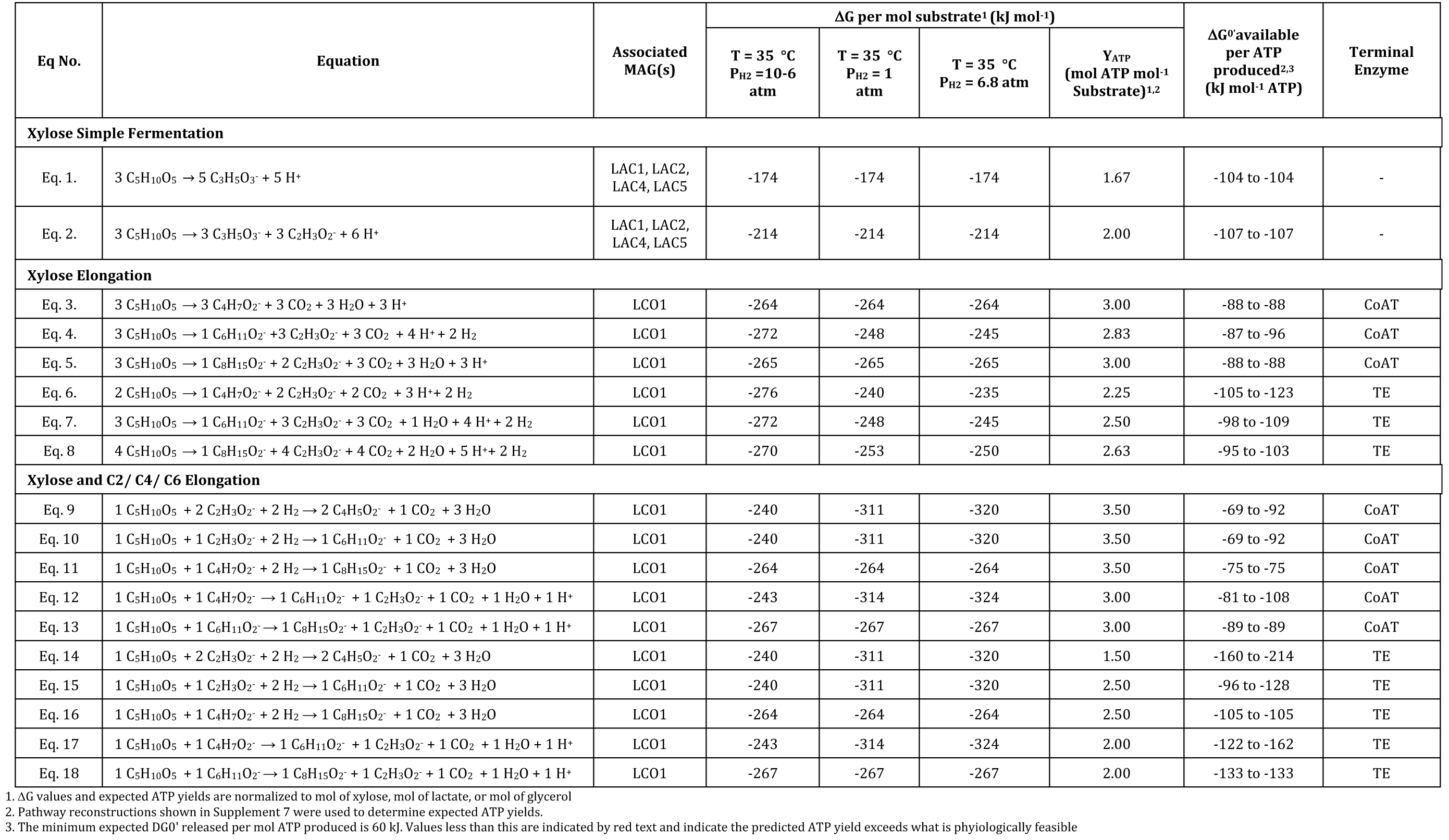
Thermodynamics of biochemical reactions involved in conversion of xylose to butyrate, hexanoate and octanoate. Free energies of formation for all chemical compounds were obtained from Kbase (www.kbase.us). The ATP yield was determined based on biochemical models presented in Supplement 7 and is indicated as mol ATP produced per mol xylose or lactate consumed. The terminal enzyme of reverse β-oxidation, either a CoA transferase (CoAT) or thioesterase (TE), is also indicated.

| Eq No. | Equation | Associated<br>MAG(s) | ΔG per mol substrate<sup>1</sup> (kJ mol<sup>-1</sup>) |  |  |  | ΔG<sup>0</sup> available<br>per ATP<br>produced<sup>2,3</sup><br>(kJ mol<sup>-1</sup> ATP) | Terminal<br>Enzyme |
| --- | --- | --- | --- | --- | --- | --- | --- | --- |
|  |  |  | T = 35 °C<br>P<sub>H2</sub> =10-6<br>atm | T = 35 °C<br>P<sub>H2</sub> = 1<br>atm | T = 35 °C<br>P<sub>H2</sub> = 6.8 atm | Y<sub>ATP</sub><br>(mol ATP mol<sup>-1</sup><br>Substrate)<sup>1,2</sup> |  |  |
| Xylose Simple Fermentation |  |  |  |  |  |  |  |  |
| Eq. 1. | 3 C<sub>5</sub>H<sub>10</sub>O<sub>5</sub> → 5 C<sub>3</sub>H<sub>5</sub>O<sub>3</sub><sup>-</sup> + 5 H<sup>+</sup> | LAC1, LAC2,<br>LAC4, LAC5 | -174 | -174 | -174 | 1.67 | -104 to -104 | - |
| Eq. 2. | 3 C<sub>5</sub>H<sub>10</sub>O<sub>5</sub> → 3 C<sub>3</sub>H<sub>5</sub>O<sub>3</sub><sup>-</sup> + 3 C<sub>2</sub>H<sub>3</sub>O<sub>2</sub><sup>-</sup> + 6 H<sup>+</sup> | LAC1, LAC2,<br>LAC4, LAC5 | -214 | -214 | -214 | 2.00 | -107 to -107 | - |
| Xylose Elongation |  |  |  |  |  |  |  |  |
| Eq. 3. | 3 C<sub>5</sub>H<sub>10</sub>O<sub>5</sub> → 3 C<sub>4</sub>H<sub>7</sub>O<sub>2</sub><sup>-</sup> + 3 CO<sub>2</sub> + 3 H<sub>2</sub>O + 3 H<sup>+</sup> | LCO1 | -264 | -264 | -264 | 3.00 | -88 to -88 | CoAT |
| Eq. 4. | 3 C<sub>5</sub>H<sub>10</sub>O<sub>5</sub> → 1 C<sub>6</sub>H<sub>11</sub>O<sub>2</sub><sup>-</sup> +3 C<sub>2</sub>H<sub>3</sub>O<sub>2</sub><sup>-</sup> + 3 CO<sub>2</sub> + 4 H<sup>+</sup> + 2 H<sub>2</sub> | LCO1 | -272 | -248 | -245 | 2.83 | -87 to -96 | CoAT |
| Eq. 5. | 3 C<sub>5</sub>H<sub>10</sub>O<sub>5</sub> → 1 C<sub>8</sub>H<sub>15</sub>O<sub>2</sub><sup>-</sup> + 2 C<sub>2</sub>H<sub>3</sub>O<sub>2</sub><sup>-</sup> + 3 CO<sub>2</sub> + 3 H<sub>2</sub>O + 3 H<sup>+</sup> | LCO1 | -265 | -265 | -265 | 3.00 | -88 to -88 | CoAT |
| Eq. 6. | 2 C<sub>5</sub>H<sub>10</sub>O<sub>5</sub> → 1 C<sub>4</sub>H<sub>7</sub>O<sub>2</sub><sup>-</sup> + 2 C<sub>2</sub>H<sub>3</sub>O<sub>2</sub><sup>-</sup> + 2 CO<sub>2</sub> + 3 H<sup>+</sup> + 2 H<sub>2</sub> | LCO1 | -276 | -240 | -235 | 2.25 | -105 to -123 | TE |
| Eq. 7. | 3 C<sub>5</sub>H<sub>10</sub>O<sub>5</sub> → 1 C<sub>6</sub>H<sub>11</sub>O<sub>2</sub><sup>-</sup> + 3 C<sub>2</sub>H<sub>3</sub>O<sub>2</sub><sup>-</sup> + 3 CO<sub>2</sub> + 1 H<sub>2</sub>O + 4 H<sup>+</sup> + 2 H<sub>2</sub> | LCO1 | -272 | -248 | -245 | 2.50 | -98 to -109 | TE |
| Eq. 8. | 4 C<sub>5</sub>H<sub>10</sub>O<sub>5</sub> → 1 C<sub>8</sub>H<sub>15</sub>O<sub>2</sub><sup>-</sup> + 4 C<sub>2</sub>H<sub>3</sub>O<sub>2</sub><sup>-</sup> + 4 CO<sub>2</sub> + 2 H<sub>2</sub>O + 5 H<sup>+</sup> + 2 H<sub>2</sub> | LCO1 | -270 | -253 | -250 | 2.63 | -95 to -103 | TE |
| Xylose and C2/ C4/ C6 Elongation |  |  |  |  |  |  |  |  |
| Eq. 9 | 1 C<sub>5</sub>H<sub>10</sub>O<sub>5</sub> + 2 C<sub>2</sub>H<sub>3</sub>O<sub>2</sub><sup>-</sup> + 2 H<sub>2</sub> → 2 C<sub>4</sub>H<sub>5</sub>O<sub>2</sub><sup>-</sup> + 1 CO<sub>2</sub> + 3 H<sub>2</sub>O | LCO1 | -240 | -311 | -320 | 3.50 | -69 to -92 | CoAT |
| Eq. 10 | 1 C<sub>5</sub>H<sub>10</sub>O<sub>5</sub> + 1 C<sub>2</sub>H<sub>3</sub>O<sub>2</sub><sup>-</sup> + 2 H<sub>2</sub> → 1 C<sub>6</sub>H<sub>11</sub>O<sub>2</sub><sup>-</sup> + 1 CO<sub>2</sub> + 3 H<sub>2</sub>O | LCO1 | -240 | -311 | -320 | 3.50 | -69 to -92 | CoAT |
| Eq. 11 | 1 C<sub>5</sub>H<sub>10</sub>O<sub>5</sub> + 1 C<sub>4</sub>H<sub>7</sub>O<sub>2</sub><sup>-</sup> + 2 H<sub>2</sub> → 1 C<sub>8</sub>H<sub>15</sub>O<sub>2</sub><sup>-</sup> + 1 CO<sub>2</sub> + 3 H<sub>2</sub>O | LCO1 | -264 | -264 | -264 | 3.50 | -75 to -75 | CoAT |
| Eq. 12 | 1 C<sub>5</sub>H<sub>10</sub>O<sub>5</sub> + 1 C<sub>4</sub>H<sub>7</sub>O<sub>2</sub><sup>-</sup> → 1 C<sub>6</sub>H<sub>11</sub>O<sub>2</sub><sup>-</sup> + 1 C<sub>2</sub>H<sub>3</sub>O<sub>2</sub><sup>-</sup> + 1 CO<sub>2</sub> + 1 H<sub>2</sub>O + 1 H<sup>+</sup> | LCO1 | -243 | -314 | -324 | 3.00 | -81 to -108 | CoAT |
| Eq. 13 | 1 C<sub>5</sub>H<sub>10</sub>O<sub>5</sub> + 1 C<sub>6</sub>H<sub>11</sub>O<sub>2</sub><sup>-</sup> → 1 C<sub>8</sub>H<sub>15</sub>O<sub>2</sub><sup>-</sup> + 1 C<sub>2</sub>H<sub>3</sub>O<sub>2</sub><sup>-</sup> + 1 CO<sub>2</sub> + 1 H<sub>2</sub>O + 1 H<sup>+</sup> | LCO1 | -267 | -267 | -267 | 3.00 | -89 to -89 | CoAT |
| Eq. 14 | 1 C<sub>5</sub>H<sub>10</sub>O<sub>5</sub> + 2 C<sub>2</sub>H<sub>3</sub>O<sub>2</sub><sup>-</sup> + 2 H<sub>2</sub> → 2 C<sub>4</sub>H<sub>5</sub>O<sub>2</sub><sup>-</sup> + 1 CO<sub>2</sub> + 3 H<sub>2</sub>O | LCO1 | -240 | -311 | -320 | 1.50 | -160 to -214 | TE |
| Eq. 15 | 1 C<sub>5</sub>H<sub>10</sub>O<sub>5</sub> + 1 C<sub>2</sub>H<sub>3</sub>O<sub>2</sub><sup>-</sup> + 2 H<sub>2</sub> → 1 C<sub>6</sub>H<sub>11</sub>O<sub>2</sub><sup>-</sup> + 1 CO<sub>2</sub> + 3 H<sub>2</sub>O | LCO1 | -240 | -311 | -320 | 2.50 | -96 to -128 | TE |
| Eq. 16 | 1 C<sub>5</sub>H<sub>10</sub>O<sub>5</sub> + 1 C<sub>4</sub>H<sub>7</sub>O<sub>2</sub><sup>-</sup> + 2 H<sub>2</sub> → 1 C<sub>8</sub>H<sub>15</sub>O<sub>2</sub><sup>-</sup> + 1 CO<sub>2</sub> + 3 H<sub>2</sub>O | LCO1 | -264 | -264 | -264 | 2.50 | -105 to -105 | TE |
| Eq. 17 | 1 C<sub>5</sub>H<sub>10</sub>O<sub>5</sub> + 1 C<sub>4</sub>H<sub>7</sub>O<sub>2</sub><sup>-</sup> → 1 C<sub>6</sub>H<sub>11</sub>O<sub>2</sub><sup>-</sup> + 1 C<sub>2</sub>H<sub>3</sub>O<sub>2</sub><sup>-</sup> + 1 CO<sub>2</sub> + 1 H<sub>2</sub>O + 1 H<sup>+</sup> | LCO1 | -243 | -314 | -324 | 2.00 | -122 to -162 | TE |
| Eq. 18 | 1 C<sub>5</sub>H<sub>10</sub>O<sub>5</sub> + 1 C<sub>6</sub>H<sub>11</sub>O<sub>2</sub><sup>-</sup> → 1 C<sub>8</sub>H<sub>15</sub>O<sub>2</sub><sup>-</sup> + 1 C<sub>2</sub>H<sub>3</sub>O<sub>2</sub><sup>-</sup> + 1 CO<sub>2</sub> + 1 H<sub>2</sub>O + 1 H<sup>+</sup> | LCO1 | -267 | -267 | -267 | 2.00 | -133 to -133 | TE |

1. AG values and expected ATP yields are normalized to mol of xylose, mol of lactate, or mol of glycerol
2. Pathway reconstructions shown in Supplement 7 were used to determine expected ATP yields.
3. The minimum expected DG<sup>0</sup> released per mol ATP produced is 60 kJ. Values less than this are indicated by red text and indicate the predicted ATP yield exceeds what is physiologically feasible

**Table S3.**
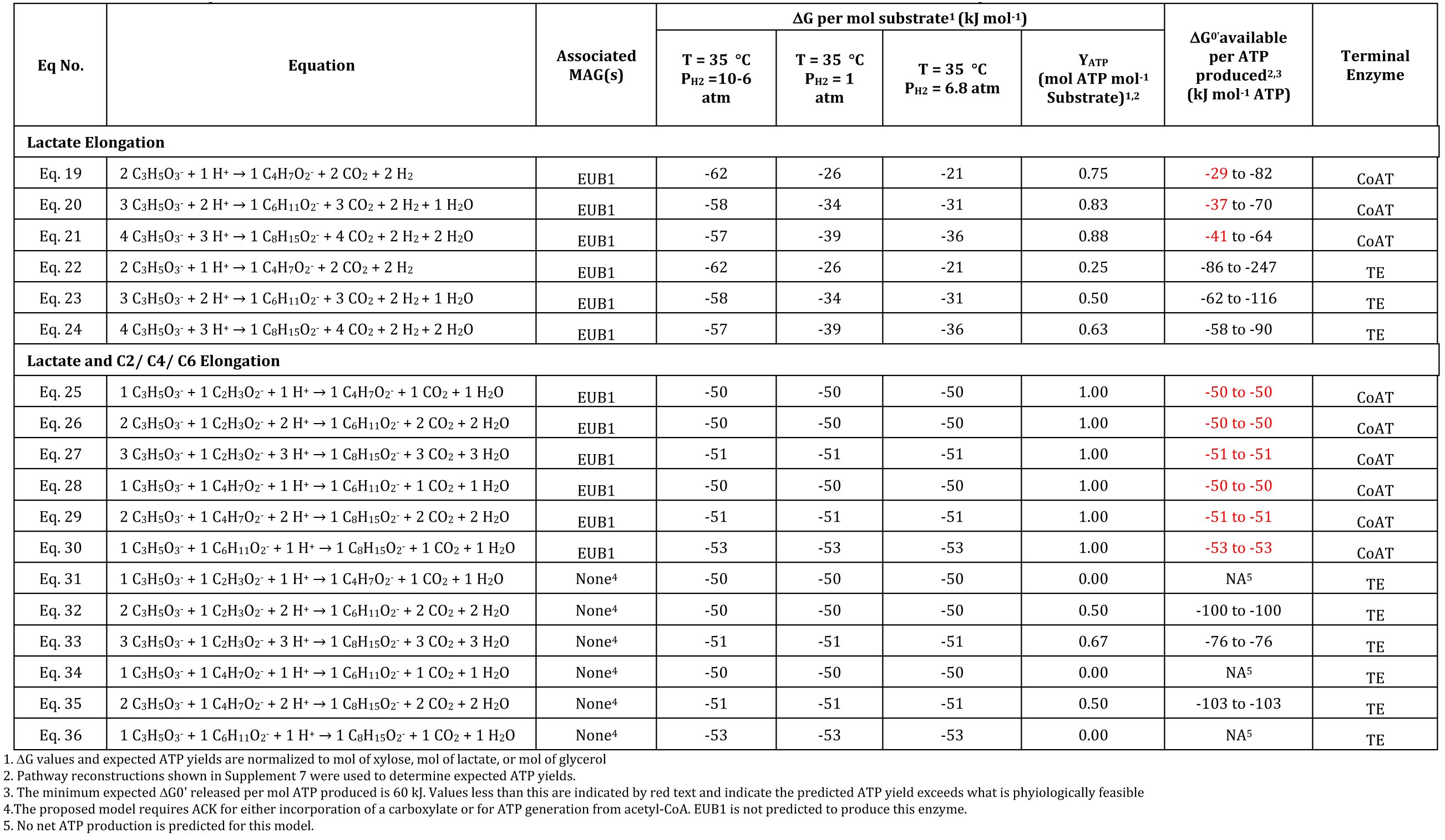
Thermodynamics of biochemical reactions involved in conversion of lactate to butyrate, hexanoate and octanoate.

